# Topographically organized dorsal raphe activity modulates forebrain sensory-motor computations and adaptive behaviors

**DOI:** 10.1101/2025.03.06.641626

**Authors:** Aytac Kadir Mutlu, Bram Serneels, Christoph Wiest, Anh-Tuan Trinh, Ricarda Bardenhewer, Fabrizio Palumbo, Oda Bjørnevik Frisvold, Inger Kristine Fjeldskaar Aukrust, Anna Maria Ostenrath, Emre Yaksi

## Abstract

The dorsal raphe nucleus (DRN) plays an important role in shaping a wide range of behaviors, including mood, motivation, appetite, sleep, and social interactions. Reflecting these diverse roles, the DRN is composed of molecularly distinct and topographically organized groups of neurons that target specific regions of the forebrain. Despite these insights, fundamental questions remain regarding how DRN neurons process sensory information, what do DRN communicate to forebrain, and the role of DRN inputs in forebrain computations and animal behavior. To address these questions, we investigated the spatiotemporal activity patterns of DRN neurons, along with DRN axons and their targets in the juvenile zebrafish forebrain. Our findings revealed a remarkable topographic organization of ongoing activity and sensory-motor responses within the DRN. We discovered that a large fraction of DRN neurons are primarily driven by animals’ locomotor activity. We also observed that an anterior group of DRN neurons, marked by Gad1, exhibited distinct activity patterns during rest, locomotor activity and sensory stimulation. DRN axons broadly innervating the forebrain exhibit topographically organized excitation and inhibition in response to sensory stimulation and motor activity. Notably, we observed significant and rapid covariation between the activity of DRN axons and nearby forebrain neurons. Chemogenetic ablation of the DRN led to a marked reduction in the synchrony and sensory-motor responses across forebrain neurons, accompanied by significant deficits in adaptive behaviors. Collectively, our findings revealed the functional diversity of DRN neurons and their role in transmitting sensory and locomotor signals via topographically organized projections, which can regulate forebrain activity and play a crucial role in modulating animal behavior.

## INTRODUCTION

Neuromodulatory systems are essential in shaping brain computations and animal behavior by dynamically regulating neural circuit activity [1-16]. Serotonin is a critical neuromodulator that is evolutionarily conserved across species [17-25].

Various serotonin receptors are widely expressed throughout the nervous system [26-28], adjusting synaptic transmission [29, 30], neural activity [16, 20, 31-33] and orchestrating network dynamics [33, 34], enabling flexible responses to changing internal states and environmental demands. The primary hub of the serotonergic system in the vertebrate brain is the dorsal raphe nucleus (DRN), located in the midbrain [19, 20, 35-37]. DRN neurons project broadly across the brain, influencing diverse neural processes from sensory and motor computations [11, 26, 38-40] to regulation of emotions and mood [4, 39, 41-43], appetite [44, 45], sleep [20, 46] as well as higher-order cognitive functions such as learning [16, 26, 38], decision-making [41, 47]. Hence, DRN is critical for the brain to correctly integrate sensory cues and adapt behavioral outputs. Consequently, DRN is a primary target for understanding and treating various neuropsychiatric disorders [48, 49].

Recent studies revealed that DRN is not a homogenous nucleus but instead composed of molecularly diverse and topographically organized groups of neurons [36, 50-53]. Viral tracing of DRN projections in mice showed that serotonergic neurons located in ventral DRN and co-expressing Vglut3 primarily innervate cortical regions [35, 54]. Whereas dorsal DRN neurons co-expressing Trh/Gad1/Gad2 mainly project to subcortical regions [35, 54]. These results suggest that molecularly and topographically distinct subcircuits of DRN neurons may be involved in processing of different neural information and distribute this to distinct forebrain targets. Do these DRN subcircuits represent functionally distinct ensembles of neurons? What kind of environmental and self-generated cues are encoded across DRN neurons? How is this information transmitted by DRN axons in the forebrain, targeting cortical and subcortical structures? And finally, what role DRN inputs play in orchestrating and regulating the activity of neural ensembles across the forebrain?

We aimed to answer these questions by using a small vertebrate, zebrafish, with relatively transparent brains that enable the monitoring of activity in large populations of individual neurons [4, 7, 19, 38, 40, 55-59] across DRN [19, 20, 26, 38, 46] and the forebrain [60, 61]. At the juvenile stage, around 3-4 weeks, zebrafish begin to exhibit cognitively demanding behaviors such as learning [62-64], social interactions [61, 65, 66], and diverse adaptive behaviors [67-69] that are typically associated with neuromodulation and maturation of the forebrain. Various developmental [70-73], molecular [70, 71], anatomical [70, 71], behavioral [74, 75] and functional [76-80] studies increasingly point toward similarities between the zebrafish and the mammalian forebrain. While most studies on the zebrafish DRN [38, 40, 81, 82] focus on reflex-like midbrain and hindbrain sensory-motor computations, the impact of DRN inputs to ancestral cortico-limbic structures in zebrafish forebrain and adaptive behaviors remains to be explored.

In this study, we examined the activity of DRN neurons and their forebrain projections in awake, behaving juvenile zebrafish. We found that DRN neurons are organized into functional ensembles, where nearby neurons exhibit correlated activity patterns during rest, sensory stimulation, and locomotion. The majority of DRN neurons are significantly modulated by the animal’s locomotor activity, while smaller subsets respond to mechanical vibrations and light. A specific ensemble of Gad1b-expressing DRN neurons, located in the anterio-dorsal DRN, displays coordinated ongoing activity and is preferentially inhibited during sensory-motor events. Imaging of DRN axons in the forebrain revealed prominent functional topography, with spatially distinct forebrain regions receiving synchronized DRN inputs that are either inhibited or excited during locomotor or sensory-evoked activity. DRN axons also exhibit strong positive and negative correlations with nearby forebrain neurons, suggesting rapid functional coupling between them. Chemogenetic DRN ablation significantly reduces synchrony across the entire forebrain, emphasizing the DRN’s role as a major orchestrator of the forebrain. Furthermore, DRN ablation decreases locomotor and sensory-evoked activation of forebrain neurons. Finally, DRN ablation significantly weakens the relationship between forebrain activity and locomotor events, impairing the animals’ recovery following aversive vibrations and their behavioral adaptation to novel tank.

## RESULTS

### Dorsal raphe nucleus is composed of topographically organized neural ensembles

DRN neurons were shown to be molecularly diverse and topographically organized [36, 50-53]. Hence, we first asked whether one can identify functional subcircuits or neuronal ensembles [83] within DRN. To do this, we performed two-photon calcium imaging of the entire DRN in head-restrained[60, 67], awake and behaving juvenile (3 weeks old) Tg(tph2:Gal4; UAS:GCaMP6s) zebrafish [40, 59, 84, 85], expressing GCaMP6s in ∼80 DRN neurons labelled by tph2 gene (Figure 1A-C, n=12). We observed a substantial level of ongoing calcium activity across the entire DRN (Figure 1D). K-means clustering [60, 86, 87] of DRN neurons based on the similarities of their ongoing activity revealed 4 optimal clusters (Figure S 1A, B). We observed that functional clusters of DRN neurons with similar activity (Figure 1D) were topographically organized into distinct DRN zones (Figure 1E). To quantify this functional DRN topography further, we plotted the average pairwise correlation of DRN neurons as a function of distance between them [59, 67, 86-88]. We observed that nearby DRN neurons exhibited more correlated ongoing activity compared to distant DRN neurons (Figure 1F). Next, we assessed the stability of DRN clusters by quantifying the likelihood that pairs of DRN neurons remain in the same cluster during two consecutive time periods, a measure we termed “cluster fidelity” [60, 86, 87]. Our analysis revealed that over 50% of DRN neuron pairs remained in the same cluster, a proportion significantly above chance levels (Figure 1G). In line with this, we also observed that correlations between DRN neurons remained stable during consecutive time periods (Figure 1H). These findings demonstrate that juvenile zebrafish DRN is composed of functionally heterogenous and topographically organized functional ensembles of neurons.

**Figure 1:**
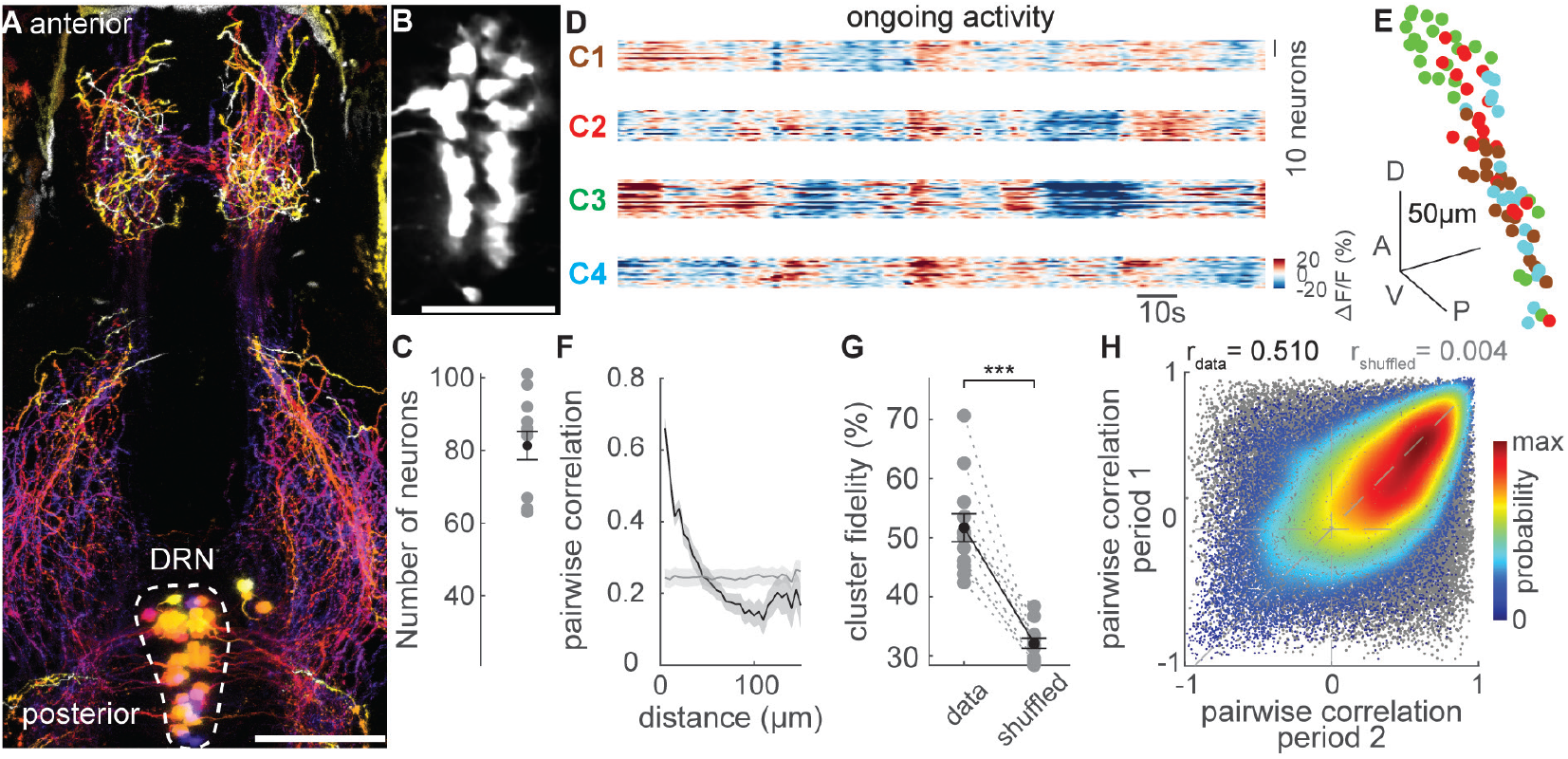
Ongoing activity of dorsal raphe ensembles is topographically organized. (A) Confocal microscopy image of dorsal raphe and its axonal projections in three-week-old Tg(tph2:Gal4;UAS:caax-GFP) juvenile zebrafish. Depth is color-coded: dorsal regions warmer and ventral regions colder. Scale bar is 100 μm. (B) Two-photon microscopy image of dorsal raphe neurons in Tg(tph2:Gal4;UAS:GCaMP6s) zebrafish. Scale bar is 50 μm. (C) 81 ± 4 (mean ± SEM) dorsal raphe neurons were recorded in each fish (n = 12). (D) Ongoing dorsal raphe activity recorded using two-photon calcium imaging in Tg(tph2:Gal4;UAS:GCaMP6s) zebrafish. Dorsal raphe ensembles are clustered (C1–4) using k-means clustering. Warm colors represent higher calcium signals. (E) Three-dimensional reconstruction of dorsal raphe ensembles (k-means functional clusters). Neurons are color-coded based on their cluster identities shown in panel D. A: anterior, P: posterior, D: dorsal, V: ventral. (F) Pairwise Pearson’s correlation of dorsal raphe neurons during ongoing activity as a function of distance (μm) between each neuron pair. Light-gray line represents shuffled spatial distribution. Shading represents SEM. (G) The ratio of dorsal raphe neuron pairs remaining in the same functional clusters during two consecutive ongoing activity periods (high cluster fidelity) is significantly higher than chance levels. (n = 12 fish). ***: p < 0.001, Wilcoxon signed-rank test. (H) Pairwise correlations of dorsal raphe neuron activity during two consecutive ongoing activity periods. pde: probability density estimate, gray dots represent pairwise correlations shuffled for pair identities. Actual data exhibit a correlation of rdata = 0.510 for the pairwise correlation across two time periods, indicating robust synchrony between pairs of neurons. Shuffled distribution: rshuffled = 0.004.

### Dorsal raphe neurons respond to locomotor and sensory signals

DRN activity is often associated with mood, emotions and internal states [3, 6, 20, 37, 38]. We asked whether DRN neurons are sensitive to relatively transient and simpler changes in animal’s locomotor activity or the sensory environment. To test this, we simultaneously measured DRN activity alongside locomotor tail-beats in head-restrained juvenile zebrafish (Figure 2A-B). We observed that DRN neurons are highly sensitive to locomotor activity. A population of DRN neurons was excited (in red), while another population was inhibited (in blue) by locomotor tail-beats (Figure 2C-D). Across all recorded fish (n=12), 26% of DRN neurons were excited, and 40% were inhibited by each tail-beat (Figure 2E-G). Three-dimensional reconstruction of locomotion-modulated DRN neurons across all fish revealed that locomotion-inhibited neurons were predominantly located at the dorsal-anterior DRN zones, while locomotion-excited neurons were positioned toward the ventral-posterior zones (Figure 2H, Figure S 1C-D). To visualize the time courses of locomotor-related DRN responses, we performed k-means clustering. This analysis revealed that DRN ensembles exhibit locomotion-evoked excitation and inhibition with diverse temporal features (Figure S1I). Interestingly, we also observed a population of DRN neurons that showed initial inhibition followed by excitation (Figure S1I, green traces).Three-dimensional reconstruction of k-means clusters of locomotor DRN responses confirmed an overall anterior-dorsal vs. posterior-ventral organization (Figure S1I), consistent with Figure 2H.To further quantify the topography of locomotion-evoked DRN responses, we calculated pairwise correlations of DRN neuron responses as a function of distance between them [67, 86-88]. This analysis revealed significantly higher response correlations between nearby DRN neurons, exceeding chance levels (Figure 2I).

**Figure 2:**
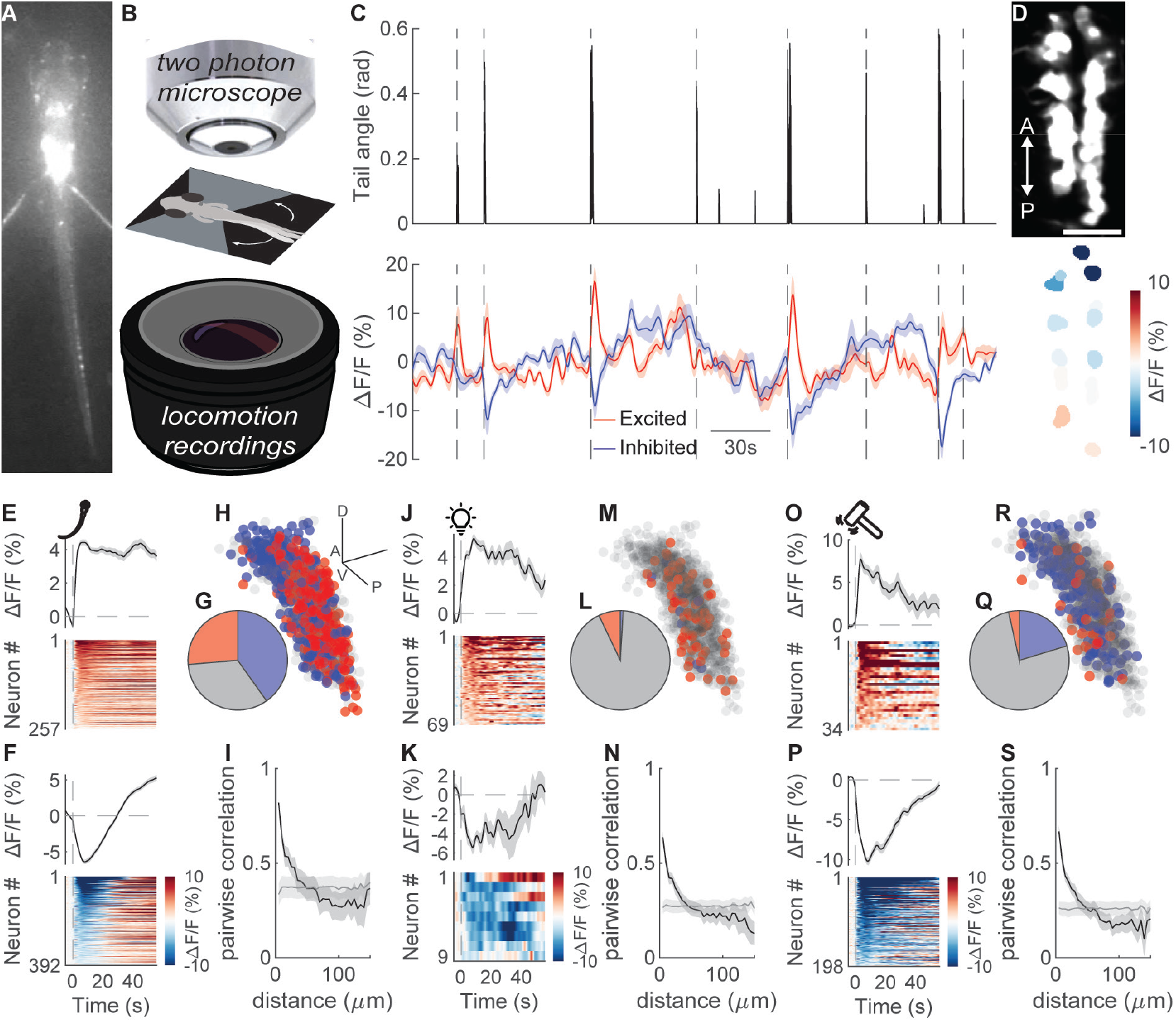
Locomotor and sensory responses of dorsal raphe neurons are topographically organized. (A) Image of a head-restrained and tail-free juvenile zebrafish recorded with an infra-red camera. (B) Scheme of the setup for simultaneous imaging of brain activity and head-restrained zebrafish locomotor behavior. (C) Example trace of locomotion measured as tail-beat angle in head-restrained juvenile zebrafish (top). Simultaneously measured calcium signals (ΔF/F) from dorsal raphe neurons (bottom). Each dashed line marks the onset of a locomotor bout. (D) Two-photon microscope image of dorsal raphe neurons in Tg(tph2:Gal4;UAS:GCaMP6s) juvenile zebrafish (top). Average neural response during the first 10 seconds of each tail-beat (bottom). Please note the inhibition (cold colors) in the anterior and the excitation (warm colors) in the posterior raphe. Scale bar represents 25 μm. (E-F) Dorsal raphe neurons excited (E, warm colors) and inhibited (F, cold colors) by locomotor tail beats. n = 12 fish. (G-H) Fraction (G) and spatial distribution (H, anterior: A, posterior: P, dorsal:D, ventral: D) of dorsal raphe neurons with significant excitation (red) and inhibition (blue) tail-beat responses. Data from all fish are spatially aligned and overlaid. Scale bar in panel H is 50 μm. (I) Pairwise Pearson’s correlation of dorsal raphe neurons as a function of their distance (μm) during tail-beat responses. Gray line represents shuffled spatial distribution. Shading represents SEM. (J-N) Same analyses as in panels E-I, during light-evoked activity (O-S) Same analyses as in panels E-I, during mechanical vibration-evoked activity Significant responses are calculated if p < 0.05 across 6 repetitions. Wilcoxon signed-rank test.

To assess sensory responses in DRN, we delivered relatively neutral red-light flashes and aversive mechanical vibrations, and performed the same analysis as described for locomotion-related signals. We observed that both red-light flashes and mechanical vibrations elicit responses in DRN. While light stimuli elicit primarily excitatory responses in 7% of DRN neurons (Figure 2J-M, Figure S1J), vibrations elicit primarily inhibition in 20% of DRN neurons (Figure 2O-R, Figure S1K). Both light and vibration stimuli, though less prominent compared to the locomotion-related activity, evoked a topographically organized DRN response based on pairwise response correlations of individual neurons (Figure 2N and S), with increased fraction vibration-evoked inhibitory responses observed towards the anterior-dorsal DRN (Figure S1H).

Altogether, these results revealed that DRN neurons are sensitive to rapid changes in animals’ locomotion and environment. Distinct zones of the DRN encode different types of locomotory and sensory information, consistent with the topographically organized DRN ensembles observed during ongoing activity (Figure 1).

### Genetically labeled dorsal raphe neurons expressing *Gad1b* exhibit distinct ongoing activity and sensory-motor responses

In rodents, a genetically distinct group of neurons co-expressing Trh/Gad1/Gad2 genes are located in the dorsal DRN and projects primarily to subcortical regions [35]. We asked whether juvenile zebrafish DRN contains such Gad1-positive neurons and whether these neurons form a distinct functional ensemble. We observed that ∼25% of DRN neurons in the juvenile zebrafish are double-labeled with Tg(Gad1b:dsRed) [89] Tg(tph2:Gal4;UAS:GCamp6s) expression (in magenta and white, Figure 3A-C). These Gad1b-labeled DRN neurons are primarily located in the dorsal-anterior zones (Figure 3A-B).

**Figure 3:**
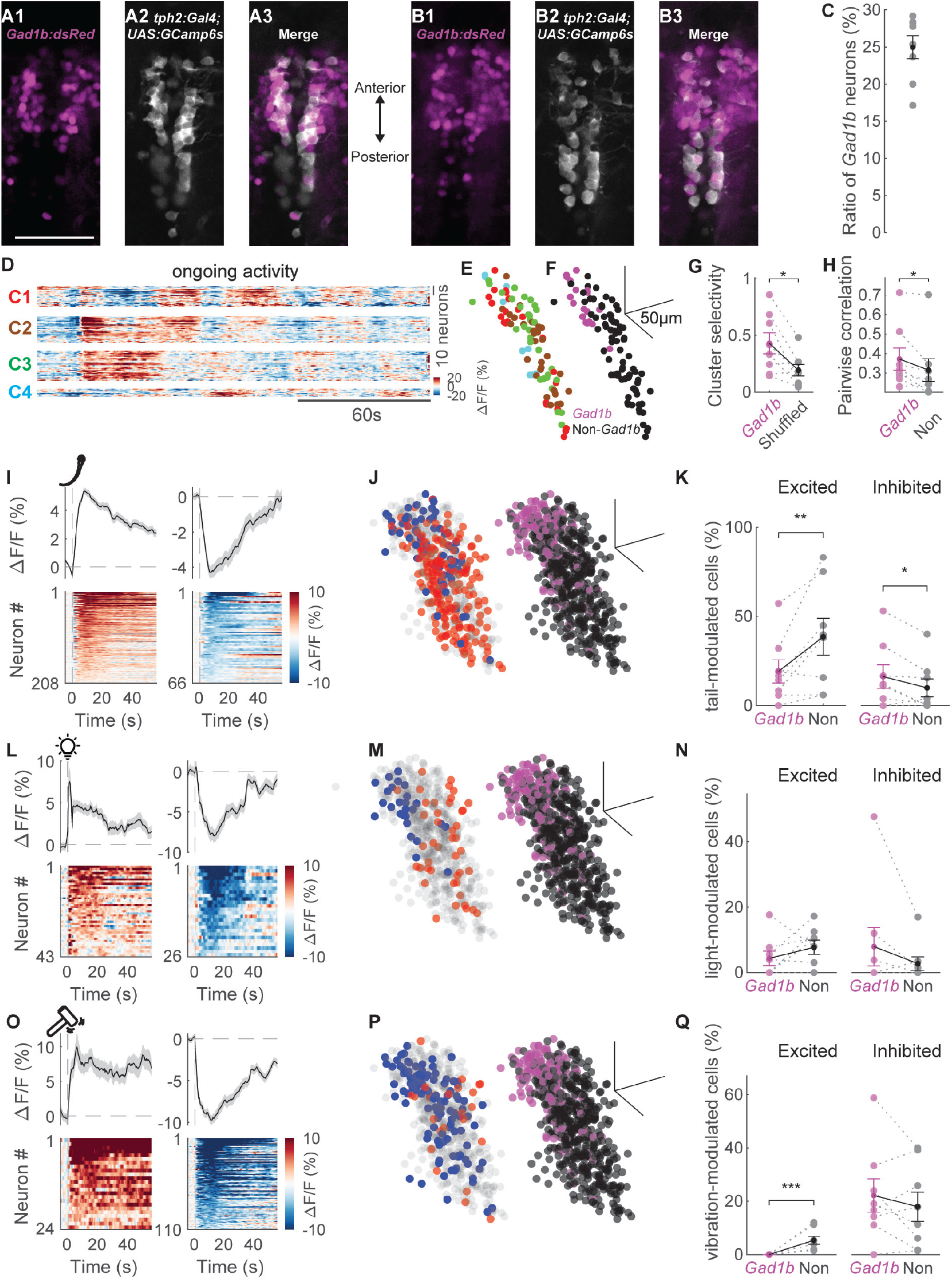
Gad1b-labeled dorsal raphe neurons exhibit distinct ongoing activity and sensory-motor responses. (A-B) Confocal microscopy images of dorsal raphe in Tg(Gad1b:dsRed ; tph2:Gal4 ; UAS:GCamp6s) juvenile zebrafish, in dorsal (A1-A3) and ventral (B1-B3) optical planes. Gad1b:dsRed expression in magenta (left), tph2:Gal4 ; UAS:GCamp6s expression in white (middle) and merged (right). (C) The ratio of Gad1b:dsRed-labeled dorsal raphe neurons. (D) Ongoing activity of dorsal raphe neurons measured by two-photon calcium imaging in Tg(Gad1b:dsRed ; tph2:Gal4 ; UAS:GCaMP6s) zebrafish. Dorsal raphe neurons are clustered (C1–4) using k-means clustering. Warm colors represent higher calcium signals. (E) Three-dimensional reconstruction of dorsal raphe functional clusters in panel D, shown in same cluster (C1-4) color codes as in panel D. (F) Gad1b:dsRed-labeling (in magenta) of same dorsal raphe neurons in panel E. A: anterior, P: posterior, D: dorsal, V: ventral. (G) Cluster selectivity of Gad1b:dsRed dorsal raphe neurons (magenta), compared to the same number of randomly selected dorsal raphe neurons (gray). Note that Gad1b:dsRed dorsal raphe neurons are significantly more selective to functional clusters. (H) Average pairwise correlations of ongoing activity between Gad1b:dsRed dorsal raphe neurons (magenta) and towards non labelled dorsal raphe neurons. Note that Gad1b:dsRed dorsal raphe neurons are significantly more correlated among themselves. (I) Tail-beat responses of excited (warm color) and inhibited (cold color) dorsal raphe neurons. Shading represents SEM. (J) Spatial distribution of dorsal raphe neurons (right) that are significantly excited (red) and inhibited (blue) by tail beats. Spatial distribution of Gad1b:dsRed dorsal raphe neurons (left). Data from all fish are spatially aligned and overlaid. (K) Fraction of Gad1b:dsRed (Gad1b), and non-Gad1b:dsRed (Non) dorsal raphe neurons that are excited (left) or inhibited (right) by tail flicks. Note that fraction of Gad1b:dsRed neurons excited and inhbited by tail-beats are significantly different than other dorsal raphe neurons. (L-N) Same analyses as in panels I-K, during light-evoked activity. (O-Q) Same analyses as in panels I-K, during mechanical vibration-evoked activity. Significant responses are calculated if p < 0.05 (Wilcoxon signed-rank test) across 6 repetitions. Scale bars are always 50μm. n = 8 fish. *: p < 0.05, Wilcoxon signed-rank test, **: p < 0.01, ***: p < 0.001, Wilcoxon signed-rank test.

To explore how Gad1b-labeled DRN neurons relate to the topographically organized functional clusters within the DRN (Figure 1-2), we simultaneously imaged genetic identity and neural activity in the red and green spectra during ongoing activity, locomotion, and sensory stimulation. During ongoing activity, we observed that individual k-means clusters of DRN ensembles (Figure 3D-E, red ensemble) largely overlap with Gad1b-labeled neurons (Figure 3F, magenta). To quantify this overlap, we used a “cluster selectivity” index [60, 86, 87]. Cluster selectivity would be “0” if Gad1b neurons were distributed equally across all DRN ensembles, and “1” if all Gad1b neurons are in a single ensemble. We observed that Gad1b-labeled DRN neurons showed significantly high cluster selectivity as compared to chance levels, suggesting they are more likely to form a functional ensemble than by chance (Figure 3G). Consistently, we also observed significantly higher correlations between Gad1b neurons than with other random DRN neurons (Figure 3H). Next, we asked whether Gad1b-labeled DRN neurons have distinct locomotor and sensory responses. We observed that a significantly smaller fraction of Gad1b neurons were excited, while a significantly higher fraction of Gad1b neurons were inhibited by locomotor tail-beats (Figure 3I-K), when compared to other DRN neurons. We also observed similar trends for sensory responses, where Gad1b neurons were largely inhibited by visual and vibration stimulation (Figure 3L-Q). Notably, no Gad1b neuron was excited by vibrations (Figure 3Q). Taken together, our results demonstrate that Gad1b-labeled DRN neurons form topographically organized ensembles with preferentially inhibitory responses to locomotor activity and sensory stimulation.

### Forebrain innervations of dorsal raphe exhibit topographically organized activity

Axonal projections of DRN neurons broadly innervate the vertebrate forebrain [35, 54, 90, 91]. However, the type of information carried via these DRN axons and the functional topography of these projections remains to be understood. To investigate this, we measured calcium signals from DRN axons across the entire forebrain of juvenile zebrafish Tg(tph2:Gal4;UAS:GCaMP6s) (Figure 4A). First, we asked how DRN axons respond to locomotor activity across multiple optical planes (Figure 4B). We observed that a prominent fraction of DRN axons in the forebrain were significantly excited or inhibited by locomotion (Figure 4C-E), consistent with DRN neuron activity (Figure 2E-G). To quantify the functional topography of these DRN projections, we calculated pairwise correlations of axonal locomotor responses as a function of the distance between them [67, 86-88]. This analysis revealed significantly higher response correlations among nearby DRN axons, exceeding chance levels (Figure 4F). To better visualize this functional topography, we spatially aligned DRN axonal signals from all recorded fish and plotted the locations of inhibitory and excitatory axonal responses to locomotion (Figure 4G). Visualizing these axonal responses as two-dimensional histograms in the dorsal (Figure 4H) and ventral (Figure 4I) telencephalon, together with the distributions in Figure 4G, revealed that spatially distinct and only partially overlapping telencephalic zones receive different locomotor-related information from the DRN. Visual and vibration-evoked activity followed similar principles but involved much smaller fractions of sensory-modulated DRN axons (Figure 4J-W). Our results suggest that the DRN communicates different types of information to distinct forebrain targets.

**Figure 4:**
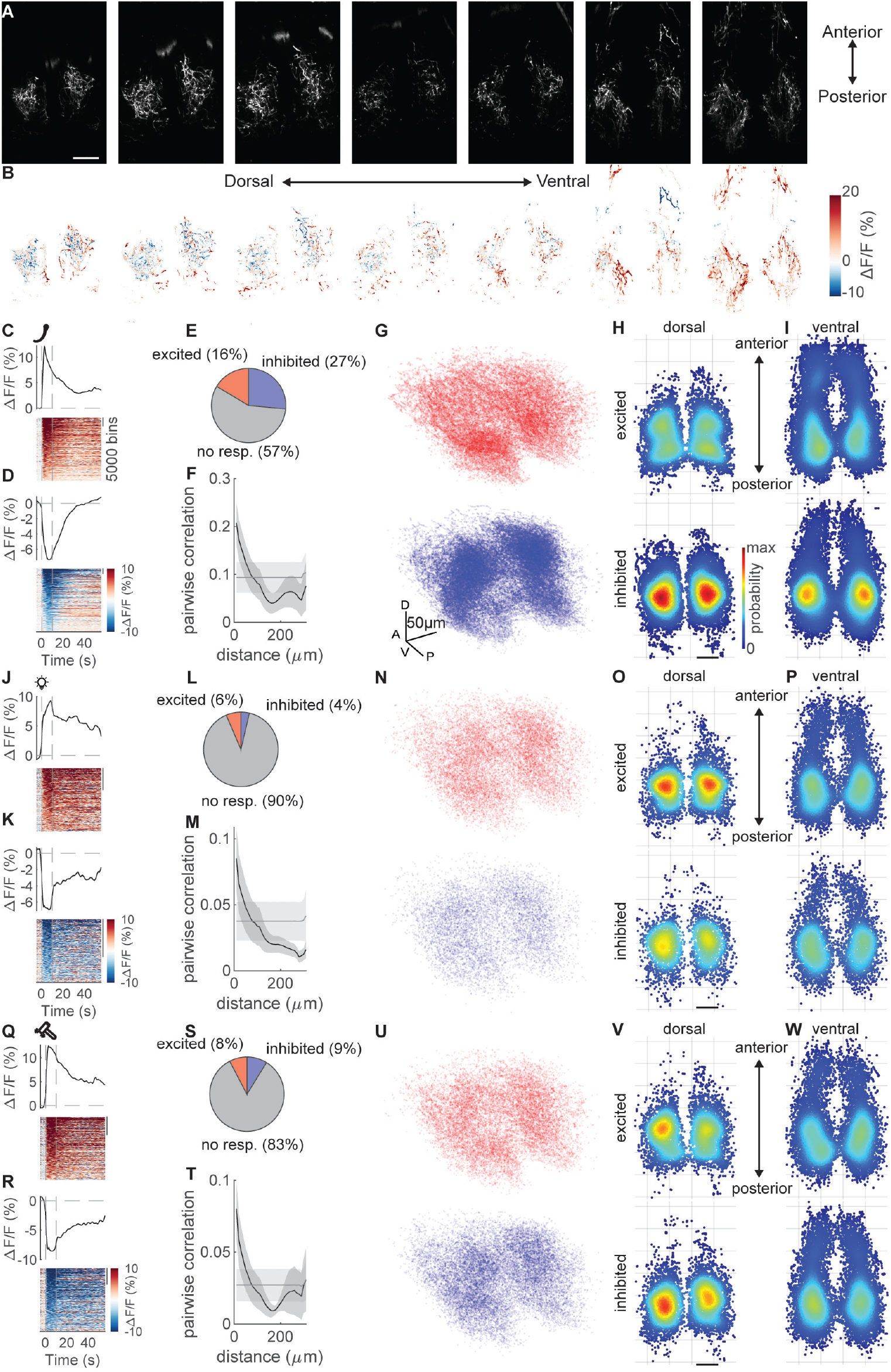
Sensory-motor responses of dorsal raphe axons innervating zebrafish forebrain are topographically organized. (A) Two-photon microscopy images of dorsal raphe axons innervating the forebrain in Tg(tph2:Gal4;UAS:GCaMP6s) juvenile zebrafish. Optical sections are shown from dorsal to ventral planes. A: anterior; P: posterior. Scale bars are 100 μm. (B) Spatial distribution of tail-beat responses (ΔF/F) of dorsal raphe axons in the forebrain. Warm colors indicate excitation, color colors indicate inhibition. Corresponding optical planes from panel A. (C-D) Tail-beat responses of excited (warm color) and inhibited (cold color) dorsal raphe axons in the forebrain. Shading represents SEM. (E) Fraction of dorsal raphe axons with significant excitation (red) and inhibition (blue) during tail-beat responses. (F) Pairwise Pearson’s correlation of dorsal raphe axons as a function of their distance (μm) during tail-beat responses. Gray line represents shuffled spatial distribution. Shading represents SEM. (G) Three-dimensional reconstruction of dorsal raphe axons that are significantly excited (red, top) and inhibited (blue, bottom) by tail beats. Significant responses are calculated if p < 0.05 across 6 repetitions of sensory stimuli and all detected locomotor tail-beats. Data from all fish are spatially aligned and overlaid. (H-I) Spatial distribution of the tail-beat excited (top) and inhibited (bottom) axonal bins in dorsal (H) and ventral (I) forebrain. Two-dimensional histograms are calculated using ks-density. Warm colors represent higher density/ probability of tail-beat modulated axons. (J-P) Same analyses as in panels C-I, during light-evoked activity. (Q-W) Same analyses as in panels C-I, during mechanical vibration-evoked activity. Scale bars are 50μm. n = 14 fish.

### Activity of the dorsal raphe projections and forebrain neuronal activity covary

We observed that DRN projections in the forebrain respond to sensory-motor stimulation with topographically organized excitation and inhibition (Figure 4). But how does this DRN axon activity relate to the activity of nearby forebrain neurons? To address this, we simultaneously imaged the activity of DRN axons together with forebrain neurons. A triple transgenic zebrafish line, Tg(tph2:Gal4; UAS:GCaMP6s; HuC:GCaMP6s-nuclear), enabled us to detect individual forebrain neuron nuclei independently from DRN axons in the forebrain, based on structural differences between axons and neuronal nuclei (Figure 5A-B; see Methods). Consistent with our results in Figure 4, we observed that mechanical vibrations elicited excitation and inhibition in different populations of DRN axons (Figure 5C-D). Next, we asked whether the activity of DRN axons in the forebrain covaries with the activity of surrounding forebrain neurons during mechanical vibrations. We observed that excited DRN axons show significantly higher positive correlations with forebrain neurons in comparison to inhibited DRN axons (Figure 5E-G).To quantify the spatial organization of these functional interactions, we calculated the correlations between the activity of DRN axons and forebrain neurons as a function of the distance between them. We observed that excited DRN axons exhibited stronger positive correlations with nearby forebrain neurons, whereas inhibited DRN axons displayed stronger negative correlations with neurons in their vicinity (Figure 5H-I).

**Figure 5:**
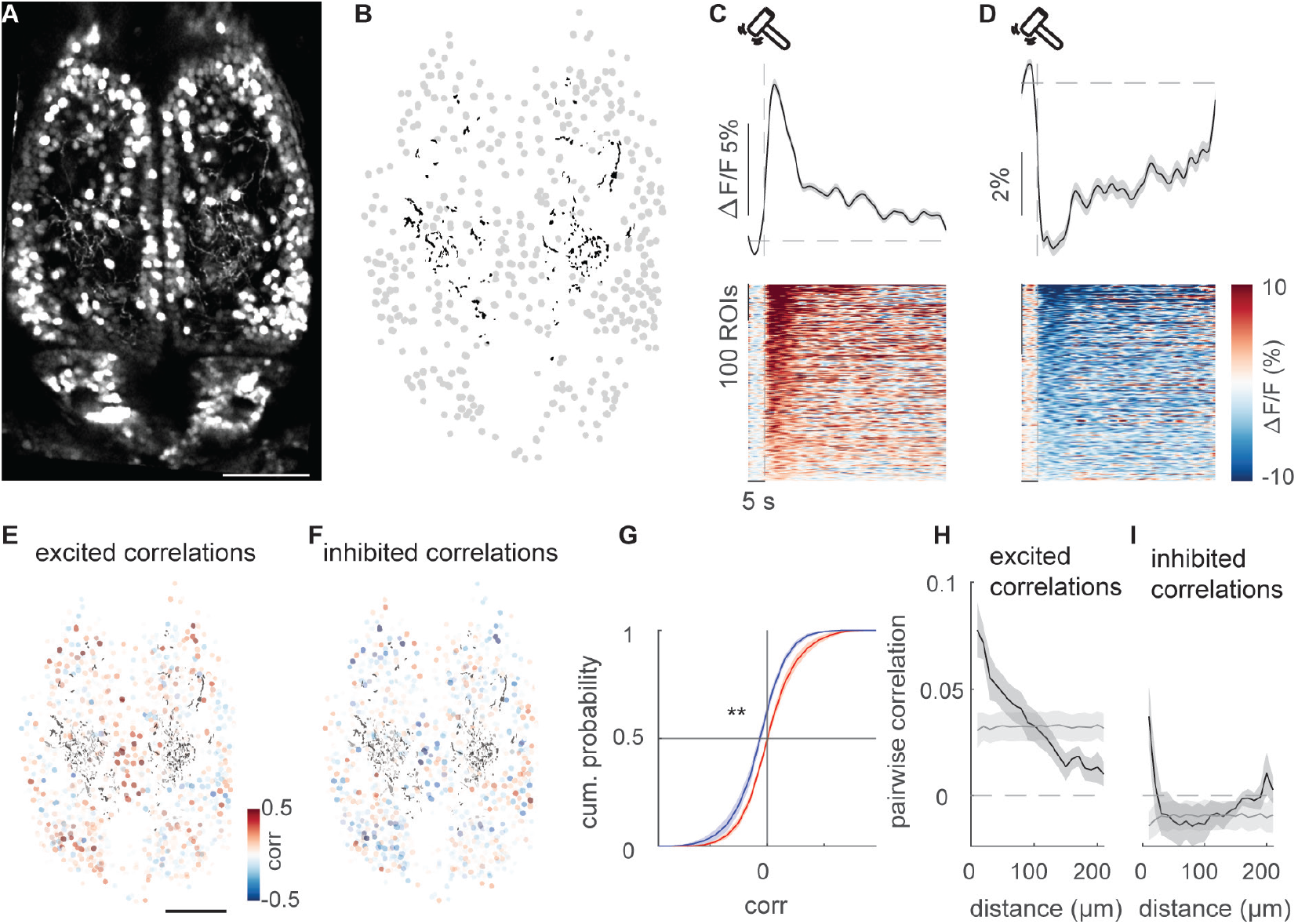
Locomotion related activity of dorsal raphe axons covaries with forebrain neurons. (A) Two-photon microscopy image of Tg(HuC:Gcamp6s-nuclear) labelled forebrain neurons together with Tg(tph2:Gal4;UAS:GCaMP6s) labelled dorsal raphe axons innervating juvenile zebrafish forebrain. (B) Two-dimensional reconstruction of dorsal raphe axons (black), and forebrain neurons (grey) identified in the image shown in panel A. (C-D) Tail-beat responses of excited (warm color) and inhibited (cold color) dorsal raphe axons in the forebrain. (E-F) Pearson’s correlations of activity between forebrain neurons and tail-beat excited (E) and inhibited (F) dorsal raphe axons. Black lines mark the dorsal raphe axons. (G) Cumulative distribution for correlations between forebrain neurons and excited (red) and inhibited (blue) dorsal raphe axons. (H-I) Pairwise Pearson’s correlation between forebrain neurons and tail-beat excited (H) and inhibited (I) dorsal raphe axonal region of interests as a function of their distance (μm) between them. Gray line represents shuffled spatial distribution. n = 8 fish. Scale bars represents 100μm. Shading represent SEM.

### Chemogenetic ablation of the dorsal raphe disrupts the synchrony and sensory-motor responses of forebrain neurons

Subsequently, we asked what role DRN projections play in regulating forebrain activity. To address this, we used a triple transgenic zebrafish expressing nitroreductase (NTR) [62, 92-94] for chemogenetic ablation of the DRN, while enabling panneuronal activity measurements, Tg(tph2:Gal4; UAS-E1b:NTR-mCherry; HuC:GCaMP6s). A 24-hour treatment with 10 mM metronidazole (MTZ) [62, 94] resulted in complete DRN ablation (Figure 6A). Given the broad forebrain innervation of DRN axons, we first asked whether DRN ablation impairs the orchestration of neurons in the dorsal zebrafish forebrain, which contains homologs of the mammalian cortico-limbic systems [70, 73, 75, 76, 95-105] and the habenula [93, 106, 107]. We quantified neural synchrony across the dorsal forebrain by calculating pairwise positive and negative correlations between neurons. We observed that both positive and negative pairwise correlations were stronger between nearby neurons and weaker between distant dorsal forebrain neurons, highlighting prominent functional topography during ongoing activity (Figure S2A, black traces) and vibration stimulation period (Figure 6B, black traces). Upon DRN ablation, both positive and negative correlations between dorsal forebrain neurons became significantly weaker (Figure 6B, Figure S2A). To test the impact of DRN ablation on the functional interactions between dorsal forebrain regions, we calculated pairwise correlations of neurons located in individual regions[108] identified using anatomical landmarks [60]. We observed an overall reduction in correlations between regions (Figure 6C, Figure S2B, cyan lines) and significant reductions in correlations between the central and posterior regions (Dm, Dc, Dmp, Dd, Hb) of the dorsal forebrain (Figure 6C, blue lines). In DRN ablated animals, neurons within individual forebrain regions also exhibited weaker positive and negative correlations that were spatially organized (Figure 6D).

**Figure 6:**
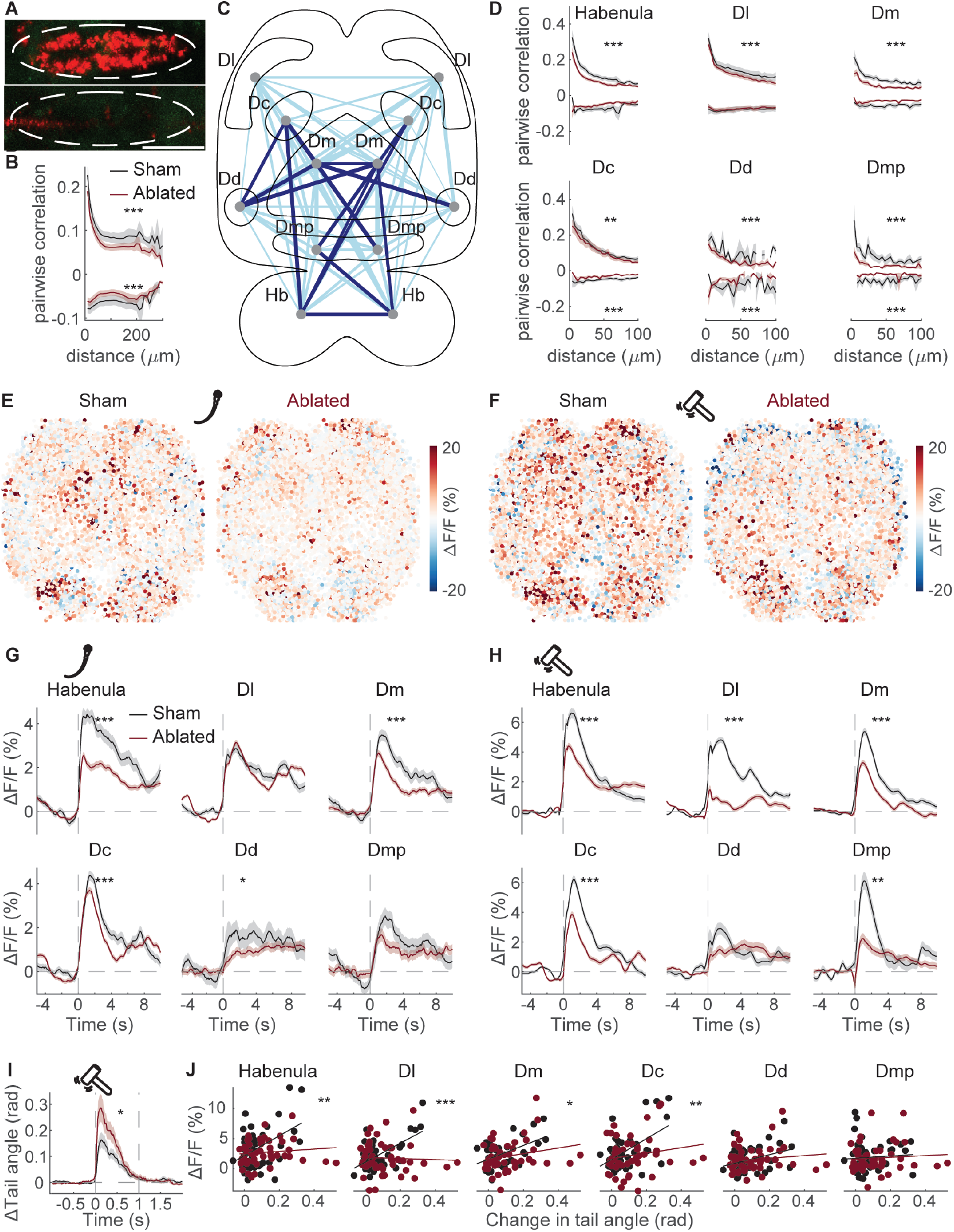
Dorsal raphe ablation impairs forebrain synchrony and sensory-motor responses. (A) Confocal microscopy image of dorsal raphe in Tg(tph2:Gal4;UAS:NTR-mCherry;HuC:GCamp6s) juvenile zebrafish after sham (DMSO, top) or Metronidazole (DMSO + MTZ, bottom) treatment. Scale bar is 50μm. (B) Pairwise positive and negative Pearson’s correlations of forebrain neurons as a function of their distance (μm) between, in sham (black, n=16 fish) and dorsal raphe ablated (brown, n=16) head-restrained juvenile zebrafish. Note an overall significant reduction of both positive and negative correlations. ANOVA displayed significance over the groups p < 0.001 is indicated with ***. (C) Schematic illustration of alterations in functional connectivity across dorsal forebrain regions upon chemogenetic dorsal raphe ablation, during vibration-evoked activity. The locations of anatomically identified dorsal forebrain regions are marked with grey dots and abbreviated. The thickness of individual lines represents the average difference in correlations between thousands of individual neurons across forebrain regions. Cyan lines represent a decrease in average correlations, and blue lines represent a significant decrease in average correlations. Dorsal raphe ablated: n=16 fish and 9709 neurons; sham: n=16 fish and 8321 neurons. Dc: Dorsal-central telencephalon, Dd: Dorsal-dorsal telencephalon, Dl: Dorsal-lateral telencephalon, Dm: Dorsal-medial telencephalon, Dmp: Dorsal-medial-posterior telencephalon, Hb: Habenula. (D) Pairwise positive and negative Pearson’s correlations of neurons within identified forebrain regions as a function of their distance (μm) between, in sham (black) and dorsal raphe ablated (brown) zebrafish. *: p < 0.05, **: p < 0.01, ***: p < 0.001 according to ANOVA. (E-F) Spatial distribution of tail-beat (E) and vibration (F) responses (ΔF/F) in dorsal forebrain neurons in sham (left) and dorsal raphe ablated (right) head-restrained juvenile zebrafish. Warm colors represent excitation, cold colors represent inhibition. Data from all fish are spatially aligned and overlaid. (G-H) Time courses of average tail-beat (G) and vibration (H) responses of neurons in individual dorsal forebrain regions in sham (black, n=16 fish) and dorsal raphe ablated (brown, n=16 fish) zebrafish. Please note the overall reduction of neural responses in dorsal raphe ablated zebrafish. Shading represents SEM. *: p < 0.05, **: p < 0.01, ***: p < 0.001, using linear mixed-effects model. # neurons in sham: Hb = 1698, Dl = 2590, Dm = 1241, Dc = 1575, Dd = 399, Dmp = 372; # neurons in ablated: Hb = 2158, Dl = 3121, Dm = 1278, Dc = 1795, Dd = 533, Dmp = 344. (I) Time courses of average behavioral responses (change in tail angle) to mechanical vibrations for sham (black), and ablated (brown) head-restrained juvenile zebrafish (same fish as in panel E-H) *: p < 0.05, using linear mixed-effects model. (J) Average behavioral response (change in tail angle) versus average neural responses in individual forebrain regions up on mechanical vibrations sham (black), and ablated (brown) zebrafish (each data point represents an individual animal). Solid lines indicate linear fits. Please note the overall reduction of relation between neural and behavioral responses in dorsal raphe ablated zebrafish. *: p < 0.05, **: p < 0.01, ***: p < 0.001 according to the likelihood-ratio test.

To further investigate the impact of the DRN on forebrain activity, we examined how forebrain responses during locomotion and aversive mechanical vibrations [67, 87] are altered by DRN perturbation. To visualize this, we spatially aligned all recorded neurons from all fish (sham: n=16 fish and 8321 neurons, ablated: n=16 and 9709 neurons) and plotted their average responses during locomotor tail-beats (Figure 6E) and vibration stimuli (Figure 6F). In DRN ablated animals, we observed weaker responses across dorsal forebrain. To determine whether responses in individual forebrain regions were differentially altered by DRN ablation, we quantified and compared response amplitudes from thousands of individual forebrain neurons. Locomotor responses were significantly weaker in central and posterior dorsal regions such as Dm, Dc, Dd, and the habenula (Figure 6G). Vibration-evoked responses were significantly weaker in Dl, Dm, Dc, Dmp, and the habenula (Figure 6H).

Our head-restrained juvenile zebrafish preparation enabled measurement of the animals’ tail-beat responses to aversive mechanical vibrations [67, 87]. Comparing the DRN-ablated group to the sham group revealed a significant increase in the amplitude of locomotor tail-beats upon mechanical vibrations (Figure 6I), while spontaneous locomotor activity during the baseline period was not changed (Figure S2C). Finally, we asked whether DRN ablation altered the relationship between neural and behavioral responses to vibrations. To address this, we compared statistical models relating the neural response of each forebrain region to animals’ behavioral responses. The null model consisted of a linear regression with the behavioral response as the sole independent variable, while the alternative model included ablation as a second independent variable (see Methods). DRN ablation significantly altered the relationship between behavioral and neural responses for the habenula, Dl, Dm, and Dc (Figure 6J). Taken together, our results highlight the importance of DRN in regulating forebrain activity and shaping its link to animals’ behavioral responses.

### Dorsal raphe ablation impairs adaptive behaviors in juvenile zebrafish

It is essential for animals to recover quickly from harmless threats, enabling them to explore their surroundings for potential resources. Such recovery requires a level of cognitive capacity to assess the intensity and history of threats, as well as to analyze associated risks and benefits. This cognitively demanding and adaptive behaviors are often attributed to cortico-limbic structures in the vertebrate forebrain [74, 75, 93]. Given our observations on role of the dorsal raphe nucleus (DRN) in regulating forebrain activity and its link to behavior (Figure 6), we set out to test the effect of DRN ablation on a set of adaptive behaviors. Juvenile and adult zebrafish are known to exhibit bottom-diving behavior in response to immediate threats [109], followed by a gradual recovery to their baseline exploratory state [67, 110-112]. We investigated the impact of DRN ablation on two such adaptive behaviors.

To do this, we quantified juvenile zebrafish behavior from a vertical view and observed that upon mechanical vibrations, juvenile zebrafish perform an immediate bottom dive, followed by a slow and gradual recovery period during which they climb upward [67] (Figure 7A, black). DRN-ablated zebrafish showed delayed upward climbing (Figure 7A), with significantly shorter distances from the bottom of the tank during the recovery period (Figure 7B). Additionally, DRN-ablated zebrafish exhibited a denser occupancy probability closer to the bottom of the tank, suggesting reduced exploration of the tank (Figure 7C). To quantify exploration, we used a focality measure, where a focality value of “1” indicates no exploration and “0” indicates complete exploration of all possible tank locations (see Methods for description). DRN-ablated zebrafish exhibited significantly higher focality, indicating reduced exploration during the recovery period (Figure 7D). When introduced to a novel tank, juvenile zebrafish display a prominent defensive behavior, performing a deep bottom dive followed by a gradual recovery period of upward climbing [113, 114] (Figure 7E). During this recovery period, DRN-ablated zebrafish exhibited significantly lower y-positions, remaining closer to the bottom of the tank compared to controls (Figure 7E-F). Furthermore, DRN-ablated zebrafish demonstrated a denser occupancy probability near the bottom of the tank, with significantly higher focality values indicating reduced exploration (Figure 7G and H). Together, these findings revealed that DRN ablated zebrafish has impaired recovery and reduced exploration after experiencing an environmental threat.

**Figure 7:**
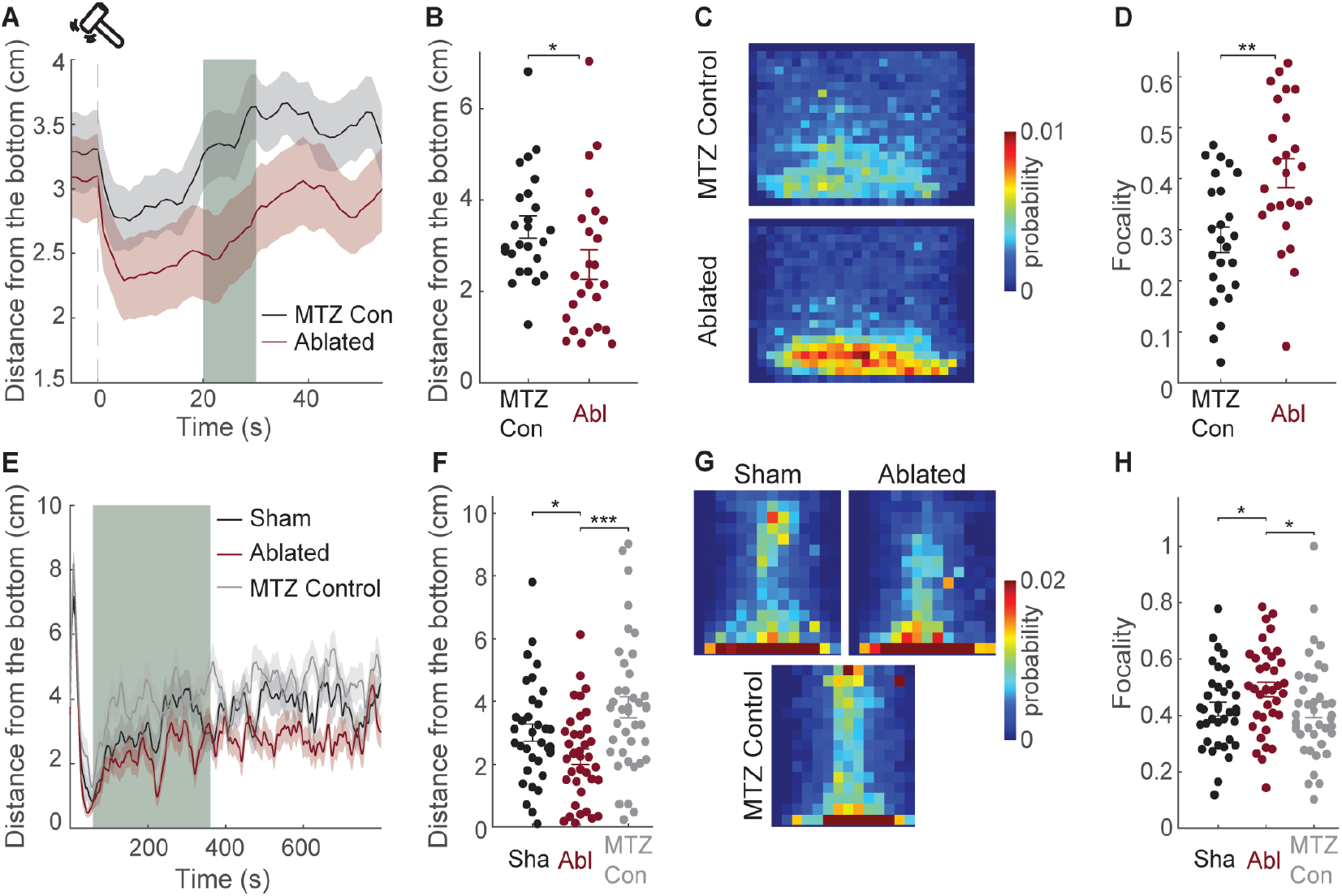
Dorsal raphe ablation impairs adaptive behaviors in freely swimming zebrafish. (A) Average time course of vertical swim position (distance from the bottom) in response to mechanical vibrations in MTZ-control (black, n = 25), and dorsal raphe ablated (brown, n = 24) juvenile zebrafish. Grey block highlights the time period that mechanical vibrations responses are quantified in panels B-D. (B) Average distance from the bottom for individual zebrafish during mechanical vibration response. Note that dorsal raphe ablated zebrafish (brown) swim significantly closer to the bottom, in comparison to MTZ-control (black) zebrafish. (C) Heatmaps representing the average position probability of freely swimming zebrafish during mechanical vibration response for the same groups as in panel A-B. (D) Focality of the position probability in panel C during mechanical vibration response. Note that dorsal raphe ablated zebrafish (brown) swim significantly more focal (less exploration), in comparison to MTZ-control (black) zebrafish. (E) Average time course of vertical swim position (distance from the bottom) during novel tank response in sham (black, n = 35), MTZ-control (grey, n = 39), and dorsal raphe ablated (brown, n = 36) juvenile zebrafish. Grey block highlights the time period that the novel tank responses are quantified in panels F-H. (F) Average distance from the bottom for individual zebrafish during novel tank response. Note that dorsal raphe ablated zebrafish (brown) swim significantly closer to the bottom, in comparison to sham (black) and MTZ-control (grey) zebrafish. (G) Heatmaps representing the average position probability of freely swimming zebrafish during the novel tank response for the same three groups as in panel E-F. (H) Focality of the position probability in panel G during the novel tank response. Note that dorsal raphe ablated zebrafish (brown) swim significantly more focal (less exploration), in comparison to sham (black) and MTZ-control (grey) zebrafish. Shadings represent SEM. (*: p < 0.05, **: p < 0.01, ***: p < 0.001, two-sided Wilcoxon ranksum test).

## DISCUSSION

### Diversity of DRN neurons and their axonal projections in the forebrain

In this study, we observed a remarkable diversity of functional features across zebrafish DRN neurons and their axonal projections in the forebrain. Mammalian DRN neurons have been shown to be diverse, with molecularly distinct groups of neurons located in topographically organized DRN zones [36, 54, 115]. Here, we revealed that the zebrafish DRN is composed of topographically organized functional ensembles, where nearby DRN neurons exhibit similar ongoing and sensory-motor-evoked activity. In vivo population recordings of individual rodent DRN neurons have recently revealed functionally distinct neural ensembles [116]. This study also suggests spatial segregation of neural ensembles recorded in a small fraction of the DRN. These findings are in line with our recordings covering the entire zebrafish DRN. Furthermore, we observed that a Gad1b-expressing population of DRN neurons exhibit functionally distinct neural activity in juvenile zebrafish. Interestingly, younger zebrafish larvae do not have such Gad1-positive DRN neurons [19, 117], suggesting a developmental increase in the molecular diversity of DRN neurons. Yet, Gad1 co-expressing neurons are present in the mice DRN [35], at a dorsal location similar to juvenile zebrafish DRN. These results indicate a conservation of developmental principles underlying the molecular topography of the DRN. Future studies on evolutionary comparison of molecularly defined DRN cell types, together with their development and topography, is needed for a better understanding of DRN function across species. In line with the functional diversity of DRN neurons, we also observed that DRN axons projecting to the zebrafish forebrain exhibit topographically organized activity. To our knowledge, functional measurements of DRN axons activity across the mammalian forebrain have not been done. Nevertheless, anatomical viral tracing experiments revealed that molecularly distinct groups of mice DRN neurons innervate distinct cortical and subcortical regions [35, 54], suggesting a specialization of DRN information received by distinct forebrain targets. The same study revealed that while Vglut3-expressing DRN neurons primarily target cortical regions, Trh/Gad1/Gad2-expressing DRN neurons target subcortical regions. We observed that locomotion-inhibited DRN axons are primarily present at a central region extending from the ventral (preoptic area of the hypothalamus, pallidum) to the dorsal (amygdala homolog Dm) zebrafish telencephalon (Figure 4G-H, in blue). Moreover, a stronger preference for locomotion-evoked inhibition in Gad1b-expressing and anterior-dorsal DRN neurons (Figure 2H, Figure 3J-K, Suppl. Figure 1D) suggests that locomotion-inhibited DRN axons may, at least in part, originate from a distinct DRN population. Indeed, Trh/Gad1 co-expressing DRN neurons in mice were shown to project to distinct hypothalamic, amygdalar, and pallidal regions [35, 54], supporting the functional specialization of forebrain-projecting DRN axons that we have observed in zebrafish. We also observed that locomotion-excited DRN axons from primarily Gad1b-negative and posterior-ventral DRN neurons (Figure 2H, Figure 3, Suppl. Figure 1C) innervate the zebrafish olfactory bulb and piriform cortex homolog Dp, as well as anterior, central, and lateral regions of the dorsal telencephalon (Figure 4G-H), which are homologous to mammalian hippocampus and cortical structures [70, 73, 75, 76, 95-105], resembling axonal innervation patterns of Vglut3-expressing DRN neurons in mice [35, 54]. Altogether, our findings provide important new evidence for differential functional features of DRN axons innervating topographically distinct forebrain regions.

### What do DRN axons communicate to the forebrain?

Recent studies provide detailed expression maps of diverse inhibitory and excitatory serotonin receptors across the mammalian [118, 119] and larval zebrafish brain [26]. These findings support the idea that DRN inputs are differentially interpreted by recipient neurons depending on their serotonin receptor profiles. Moreover, DRN neurons were shown to co-release serotonin together with GABA [120] or glutamate [121], which means DRN axons can interact with their targets through multiple mechanisms. We observed that DRN axons exhibit stronger positive correlations with nearby forebrain neurons (Figure 5H-I). A recent study showed that optogenetic activation of DRN neurons leads to a rapid increase in ongoing bursting activity in the zebrafish dorsal telencephalon [85]. These results are in line with our findings that genetic ablation of DRN leads to an overall reduction in sensory and motor responses and decorrelation of activity across the entire zebrafish dorsal telencephalon (Figure 6). Hence, the net effect of DRN inputs to the dorsal telencephalon appears to be facilitating and orchestrating.

Traditionally, DRN activity is associated with the encoding of emotions, mood [4, 39, 41-43], appetite [44, 45], sleep [20, 46], or even socialization [37, 122]. Zebrafish DRN was shown to encode alterations of ambient light levels [39]. Intriguingly, our results revealed that locomotor tail-beats and mechanical vibrations are prominent drivers of excitation and inhibition in distinct populations of zebrafish DRN neurons. This means that DRN has access to information related to rapid environmental changes and animals’ motor activity. In fact, we observed that chemogenetic DRN ablation led to a decoupling between zebrafish forebrain activity and locomotion (Figure 6J). Hence, our results suggest that DRN might be an additional pathway, in parallel to other channels [14, 123-127], communicating information about animal locomotion to the forebrain. Sensory-motor information was shown to be available also to mammalian DRN [128], suggesting that not only the anatomy [16], but also the function of DRN is conserved across the vertebrates.

### The Role of DRN in regulating brain and behavior

We propose that locomotor signals from DRN can be utilized to amplify neural and behavioral responses to sensory cues such as mechanical vibrations, as we observed in our experiments (Figure 6F-H and Figure 7A-D). This can be particularly important during adaptive behaviors, where animals make risk-benefit analyses. In fact, we observed that ablating DRN delays the initiation of exploratory behaviors after a transient threat. In line with our findings, 2 recent brain-wide imaging studies in larval zebrafish identified DRN as the primary region associated with the switch between animals’ exploration and exploitation states [129], as well as the switch between active to passive coping states[130]. Similarly, DRN was also shown to be important for motor adaptation in zebrafish larvae, during which animals have to track the outcome of their actions and adjust the gain of their locomotory activity [38]. Given all these, it is possible that the well-established role of DRN in mood and internal states [4, 20, 39, 41-46] might have evolved from humbler evolutionary beginnings, in which DRN-forebrain interactions mediate the communication between animals’ locomotor activity and sensory information processing. We therefore propose that future studies should focus not only on the role of DRN in encoding and regulating complex high-level behaviors but also on how simpler sensory-motor actions can influence such computations.

## ACKNOWLEDGEMENTS

We thank M. Ahrens (HHMI, Janelia Farm, USA), K. Kawakam (NIG, Japan), S. Higashijima (Okazaki Institute for Integrative Bioscience, Japan) and H. Burgess (NIH, USA) for transgenic lines. We thank S. Eggen, F. Acuña-Hinrichsen, V. Nguyen and our fish facility support team for technical assistance. We thank the Yaksi lab members for stimulating discussions. We thank Gil Costa for the zebrafish illustration shared open-access at scidraw.io. This work was funded by an RSO grant from Norwegian University of Science and Technology (E.Y, A.M.O.), MSCA Postdoctoral Fellowship (A-T.T.), NFR FRIPRO research grant 314212(E.Y.) and RCN Centres of Excellence scheme, project number 332640. Work in the E.Y. laboratory is funded by the Kavli Institute for Systems Neuroscience at Norwegian University of Science and Technology.

## AUTHOR CONTRIBUTIONS

Conceptualization, C.W., A.K.M., E.Y.; Methodology and data, A.K.M., B.S., R.B., P.W.S., F.P., A-T.T., E.Y.; Data Analysis, A.K.M., R.B., P.W.S., F.P., E.Y.; Investigation, all authors; Writing, A.K.M., E.Y.; Review & Editing, all authors; Funding Acquisition and Supervision, E.Y.

## INCLUSION AND ETHICS STATEMENT

Our team includes researchers from diverse origins and backgrounds. One or more of the authors of this paper self-identifies as an underrepresented minority in science.

All experimental procedures performed on zebrafish larvae and juveniles were in accordance with the Directive 2010/63/EU of the European Parliament and the Council of the European Union and approved by the Norwegian Food Safety Authorities. Animals from both sexes were used in this study.

## MATERIALS AND METHODS

### Lead contact

Further information and requests for resources and reagents should be directed to and will be fulfilled by the lead contact, Emre Yaksi

### Materials availability

This study did not generate new unique reagents.

### Data and code availability

Calcium imaging data reported in this paper will be deposited to resource center before publication. Main associated codes that are used to make figures are available at: https://github.com/yaksilab/drn_paper.

### Fish husbandry

NFSA (Norwegian Food Safety Authority) has approved the animal facility and fish maintenance. Fish were kept in 3,5L tanks in a Tecniplast ZebTec Multilinking System. Constant conditions were maintained: 28.5°C, pH 7.2, 700μSiemens. 14:10 hour light/dark cycle was preserved. Dry food (SDS100 up to 14dpf and SDS 400 for adult animals, Tecnilab BMI, the Netherlands) was given to fish twice a day, in addition to Artemia nauplii (Grade 0, Platinum Label, Argent Laboratories, Redmond, USA) once a day. From fertilization to 3dpf (days post fertilization) larvae were kept in a Petri dish with egg water (1.2g marine salt in 20L RO water, 1:1000 0.1% methylene blue) and between 3 and 5dpf in artificial fish water (AFW: 1.2 g marine salt in 20L RO water). Juvenile (3 to 4-week-old) zebrafish were used for the experiments.

Tg(tph2:Gal4;UAS:GCaMP6s), Tg(tph2:Gal4;UAS:GCamp6s;Gad1:dsRed), Tg(tph2:Gal4;UAS:Gcamp6s//HuC:Gcamp6s-nuclear), and Tg(tph2:Gal4;UAS:NTR-mCherry;HuC:GCamp6s) zebrafish lines were used for calcium imaging.

### Two-photon calcium imaging and sensory stimulation in head-restrained juvenile zebrafish

For in vivo imaging, 21 dpf juvenile fish were embedded in 2.5% low-melting-point agarose (LMP, Fisher Scientific) in the lid of a 35 mm Petri dish (FALCON). The constant perfusion of AFW buffered with 10mM NaHCO3 and bubbled with carbogen (95% O2 and 5% CO2) at 27oC was maintained during the experiment. After solidifying for 20 minutes, LMP agarose was carefully removed in front of the mouth and posterior to the swim bladder.

A two-photon microscope (Scientifica Inc.) with a 16x water immersion objective (Nikon, NA 0.8, LWD 3.0) was used for calcium imaging. For excitation, a mode-locked Ti:Sapphire laser (MaiTai Spectra-Physics) was tuned to 920 nm. Recordings were performed as either single-plane or volumetric recordings (8 planes with a Piezo (Physik Instrumente (PI))). The acquisition rate was 18.61 Hz for single-plane recordings (image size 1536×850 pixels) and 2.33 Hz per plane for volumetric scans (image size 1536×850 pixels).

First, spontaneous activity was measured for 3 minutes. Afterwards, six repetitions of sensory stimuli (red light flash or vibrations) were applied. All our sensory stimulation parameters are selected based on our earlier studies [67, 87, 134]. For the light stimulus, we used a red LED (LZ1-00R105, LedEngin; 625-nm wavelength) placed in front of the recording chamber near the tube. The light stimulus was a flash of 200 ms duration with an intensity of 0.318 mW. Vibrations were delivered via a solenoid tapper (SparkFun Electronics, ROB-10391) with a 200 ms application of 12 V. The total duration of the recordings was 30 minutes.

### Imaging of head-restrained zebrafish

For simultaneous imaging of calcium signals and locomotor tailbeats, a Manta camera (G-031, Allied Vision; recording at 120 fps) was used. A custom-built IR light source (780 nm LEDs) was placed around the microscope objective above the fish to provide maximal IR illumination.

Measuring behaviors in freely-swimming zebrafish:

For tracking behaivors in freely-swimming 21 dpf juvenile zebrafish, we used Zantiks LT set up with 6 tanks (10 cm x 11.5 cm x 3 cm) enabling experiments with six fish simultaneously while recording their vertical and horizontal positions in tanks. Ambient white light was available during all behavioral experiment. During novel tank diving experiment, the animals were gently placed into their respective tanks filled with AFW. Their movement during the experimental procedure was tracked at 15 Hz, in dimensions viewed from the side. Fish with tracking errors were identified manually and were not included in further analysis.

During vibration stimulation experiments 6 plastic arenas (11.5 cm x 11.5 cm x 1.5 cm) glued to an acrylic plate using epoxy resin were used. For analyzing the fish’s behavior, the fish were tracked at 10Hz. To deliver the vibration stimuli, a “Solenoid-tapper”-device (Solenoid 36V, Sparkfun Electronics) was coupled by a microcontroller Arduino Due to the arenas and programmed with a custom-software. After baseline recording (30 Min) without any external stimuli, the tapper began to tap on the arenas (1/60 Hz, 6 trials). All experiments were performed at room temperature around 24oC.

### Quantification and Statistical Analysis

Two-photon microscopy images were aligned using a method described in [59, 60, 132] (occasional XY drift was corrected, based on “hierarchical model-based motion estimation” [135]) or using suite2p [133]. Recordings were visually inspected for remaining motion and Z-drift, recordings with remaining motion artifacts were discarded. Regions of interest (ROIs) corresponding to neurons were automatically detected using a template matching algorithm [59, 60, 86], and visually confirmed. To calculate the time course of each neuron, pixels belonging to each ROI were averaged over time. For each ROI, fractional change in fluorescence (ΔF/F) relative to baseline was calculated.

Functional clusters of neurons were calculated by using k-means clustering algorithm in MATLAB (Figure 1D, Figure 4D) [86]. To identify optimal number of k-means clusters in the dorsal raphe, we used elbow method [136]. First, the distance of each cluster element to the centroid of that specific cluster was calculated, and this value was normalized by distances from each cluster element to every centroid. This operation is iterated for 1-30 number of cluster number ‘ks’ (Figure S 1A, black traces), and compared to shuffled/simulated data with same mean and variance, but no clustering (Figure S 1A, gray traces), and subtracted to find the peak distance between real data and simulated data with no clusters, to reveal the optimal clusters number (Figure S 1B). These analyses revealed that 3 minutes of ongoing DRN activity can be optimally represented by 4-5, which we choose to use in our analysis.

Cluster fidelity was calculated by measuring the probability of pairs of neurons being in the same cluster during two different 1.5 minute periods [86]. We compared the cluster fidelity of real k-means clusters with shuffled the cluster identities of same neurons.

Across this paper, excitation or inhibition (upon sensory or motor stimulation) was determined by comparing the average activity during baseline (5 s-period before stimulus onset) with the average activity during response window (5 s- or 10 s- or 20 s-period after stimulus onset) using one-tailed Wilcoxon signed-rank test, across 5-6 repetition of each stimuli.

In Figure 3, cluster selectivity was calculated to quantify the overlap of Gad1b-positive neurons with functional clusters of neurons identified using k-means clustering based on their spontaneous activity. Cluster selectivity index is the life-time sparseness [86] for the distribution of Gad1b-positive neurons across 4 k-means clusters. If cluster selectivity is 1, it means that Gad1b-positive neurons belong to one functional cluster. If cluster selectivity is 0, all Gad1b-positive neurons are equally distributed into all functional k-means clusters.

In Figure 4, axonal pixels were identified through thresholding. Per imaging plane, first, contrast of the average image was adjusted, then pixels with relative intensity value larger than %10 of the maximum were identified as axonal pixels.

In Figure 5, to identify axonal ROIs in raw images, first an ROI excluding pixels with cell nuclei was manually drawn. Then, an independent component analysis was performed on the fluorescence stacks [137, 138]. The resulting independent components were then manually inspected and verified. This procedure yields an axonal ROIs containing clusters of pixels with co-varying fluorescence changes.

In Figure 6, delineation of brain regions in the telencephalon was done manually on the raw image of the brain based on anatomical landmarks [139] described in previous studies in zebrafish and other teleost fish [71, 72, 99, 140, 141]. In each animal, time course of the tail angle change (TAC) in each trial was calculated using a baseline period of 1 s before the stimulus onset. Response window for calculating average TAC was 1 s. The linear model for the relationship between neural and behavioral response is given by the equation:

y_i=β_0+β_1*a_i+β_2*x_i+β_3*a_i*x_i+ε_i

y: neural response (average ΔF/F in response window),

i: animal index,

β0-3: effects,

a: dummy variable for ablation,

x: behavioral response (average TAC in response window),

ε: error term.

## SUPPLEMENTAL FIGURES

**Supplemental Figure 1**

**Figure S 1:**
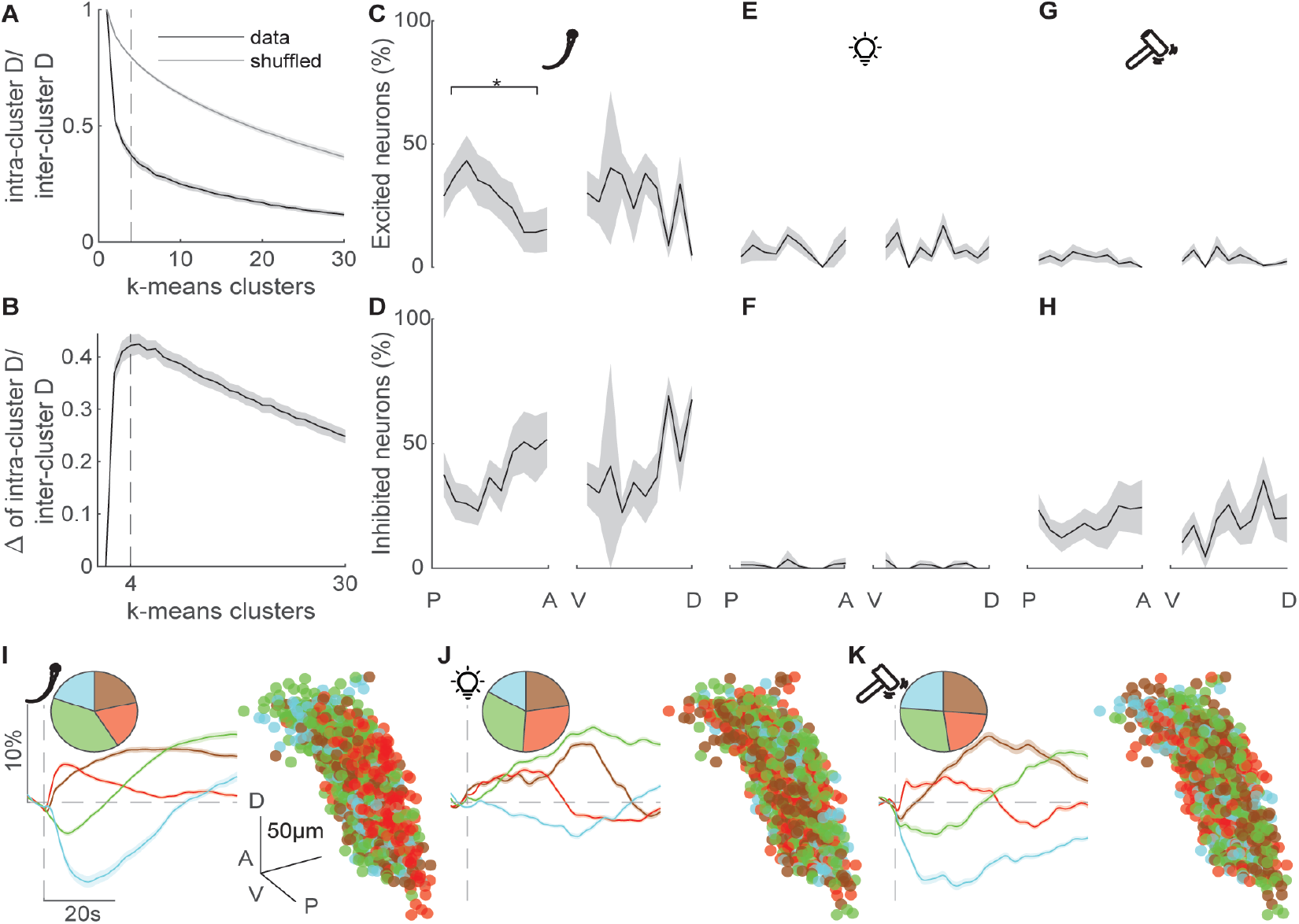
Dorsal raphe exhibits structured ongoing activity and responds to multiple stimulus modalities. Related to Figure 1 and Figure 2. ((A) Identification of optimal number of k-means clusters of ongoing dorsal raphe activity, by using elbow analysis. Elbow analysis calculates the sum of intra-cluster distances “D” for each cluster element, normalized by the sum of average inter-cluster distances for actual data (black), and simulated data (gray,100 iterations) with the same variance as the actual data but no cluster structure, in n = 12 zebrafish. This calculation is repeated for up to 30 k-means clusters (x-axis). The optimal number of clusters corresponds to the elbow point, where the black curve exhibits a prominent bend. Dashed lines indicate k-means analysis for 4 clusters. Shading represents the standard error of the mean. (B) Optimal number of clusters is further revealed, when actual data is compared to simulated data with similar variance but no cluster structure, by taking the difference of two curves in panel A. Note that the peak point of this difference reveals 4-5 optimal number of clusters in ongoing dorsal raphe activity. In this paper, we chose 4 clusters for k-means analysis of dorsal raphe activity. (C-D) Fraction of significantly excited (C) and inhibited (D) dorsal raphe neurons upon locomotor tail-beats with respect to their anteroposterior, and dorsoventral locations. A: Anterior, P: Posterior, D: Dorsal, V: Ventral. Shading represents SEM. Averages of the two most anterior/dorsal, and two most posterior/ventral location bins were compared. *: p < 0.05, Wilcoxon signed-rank test. (E-F) Same analyses as in panels C-D during light responses. (G-H) Same analyses as in panels C-D during vibration responses. (I) Responses of dorsal raphe neurons in all fish (n=12 fish, n=976 neurons), during locomotor tail-beats clustered by k-means clustering. The pie chart (top-left) represents the fraction of neurons in each color-coded k-means clusters. Time course of average responses (bottom-left) in each dorsal raphe k-means clusters. Shades represents SEM. Spatial distribution (right) of dorsal raphe neurons that are in different color-coded k-means clusters. Data from all fish are spatially aligned and overlaid. Scale bar is 50μm. (J-K) Same analyses in panel I, during light (J) and vibration (K) responses.

**Supplemental Figure 2**

**Figure S 2:**
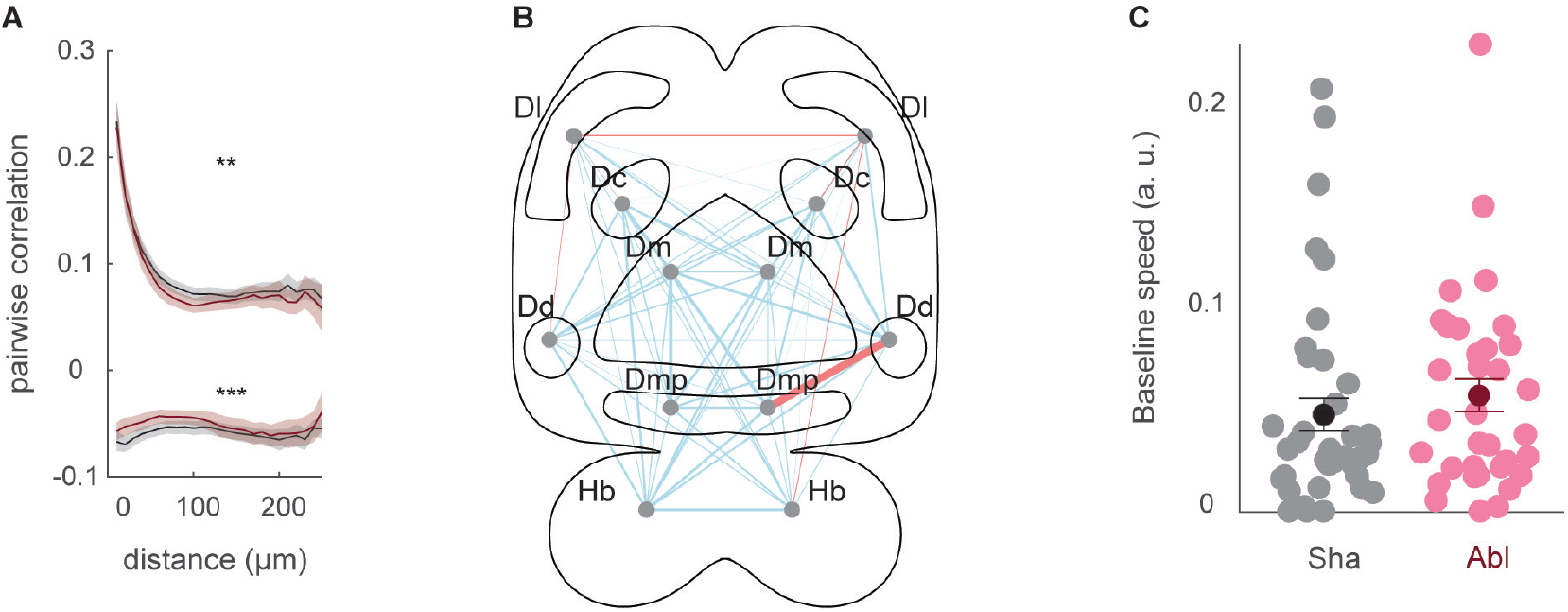
Ablation of dorsal raphe impairs ongoing forebrain synchrony, and does not affect spontaneous swimming speed in head-restrained juvenile zebrafish. Related to Figure 6. (A) Pairwise positive and negative Pearson’s correlations of forebrain neurons as a function of their distance (μm) between, in sham (black, n=16 fish) and dorsal raphe ablated (brown, n=16) head-restrained juvenile zebrafish. Note an overall significant reduction of both positive and negative correlations. ANOVA displayed significance over the groups, p < 0.01 indicated with **, p < 0.001 indicated with ***. (B) Schematic illustration of alterations in functional connectivity across dorsal forebrain regions upon chemogenetic dorsal raphe ablation, during vibration-evoked activity. The locations of anatomically identified dorsal forebrain regions are marked with grey dots and abbreviated. The thickness of individual lines represents the average difference in correlations between thousands of individual neurons across forebrain regions. Cyan lines represent a decrease in average correlations, and red lines represent an increase in average correlations. Dorsal raphe ablated: n=16 fish and 9709 neurons; sham: n=16 fish and 8321 neurons. Dc: Dorsal-central telencephalon, Dd: Dorsal-dorsal telencephalon, Dl: Dorsal-lateral telencephalon, Dm: Dorsal-medial telencephalon, Dmp: Dorsal-medial-posterior telencephalon, Hb: Habenula. (C) Average locomotion speed during the baseline period for sham (black, n = 40), and ablated (brown, n = 35) zebrafish.

**Supplemental Figure 3**

**Figure S 3:**
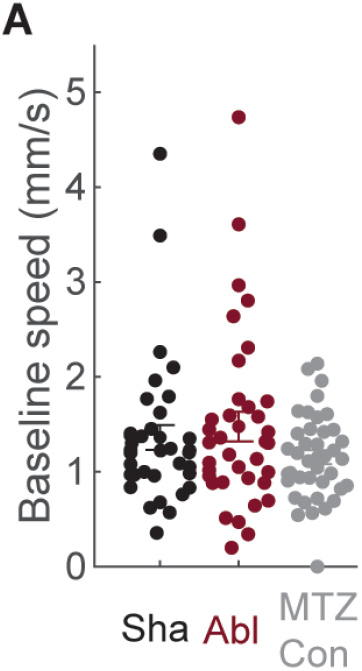
Dorsal raphe ablation does not affect free swimming speed. Related to Figure 7. (A) Average free swimming speed during baseline for sham (black, n = 35), ablated (brown, n = 36), and MTZ-treatment control (grey, n = 39) animals in the novel tank test.

